# Evaluating the Potential Blood Coagulant Activity of *Caenothus american*us Compounds: Computational Analysis using Docking, Physicochemical, and ADMET Studies

**DOI:** 10.1101/2023.09.05.555050

**Authors:** Zahra Sadat Mashkani, Zahra Pahlavan Yali, Akbar Dorgalaleh, Mahmood Shams

## Abstract

*Caenothus americanus* is a common folk remedy for the treatment of wound bleeding. Certain compounds found in this plant have been shown to reduce clotting time. However, analyzing the effects of various compounds of a folk remedy is a time-consuming and expensive process, therefore, this study employed computational analyses using docking, physicochemical, and pharmacokinetic servers to identify potential clotting activity in C.americanus compounds. The ADMETlab, SwissADME web servers, Discovery Studio, and Autodock were used to study the proper binding to target proteins and predict the physicochemical and ADMET properties (adsorption, distribution, metabolism, excretion, and toxicity) of C. americanus compounds. Coagulation factors including activated factor (F) IIa, FVa, FVIIa, FVIIIa, FIXa, FXa, FXIa, FXIIa, and FXIIIa were chosen as target proteins. Docking studies revealed that malic acid, malonic acid, oxalic acid, and succinic acid were effective on coagulation factors, of which, malic acid had better binding to intrinsic pathway factors including FVIIa, FIIa, and FXIIIa (except FVIIIa), oxalic acid to FVIIIa, and malonic acid to FVa and FXa. Moreover, ADMET studies showed the safety profile of these compounds. In conclusion, carboxylic and alcoholic groups of malic acid, malonic acid, oxalic acid and succinic acid play a role in interaction with blood coagulation factors. Additionally, based on the ADMET characteristics and suitable pharmacokinetic potentials of these compounds, they can be introduced as blood coagulant candidates with fewer side effects in bleeding disorders. However, further studies are necessary to evaluate the precise components of the *C. americanus* with the suability to bind coagulation factors.

## Introduction

Under the physiologic conditions, the coagulation system remains inactive, but once the blood vessels are damaged, it is activated to form a clot and prevent bleeding (1). Coagulation factors disorders and imbalance in this system can result in hemorrhagic and thrombotic complications (2–7). In this regard, while several therapeutic options including plasma replacement therapy, recombinant agents, gene therapy, cell therapy, and liver transplantation are available, they often come with side effects such as inhibitor production, short half-life, high costs, the risk of thrombosis (8–18), and allergic reactions (19).

Interestingly, it has been shown that medicinal plants have been one of the more important options for treating of various diseases throughout history. In this setting, active ingredients of herbal medicines been employed as a modern treatment methods in bleeding disorders (8, 9, 15, 20–28) due to their similarity to in vivo compounds (29–31). As an example of these natural compounds, there are plant phytoestrogens (32), are similar to 17-b estradiol and estrogens in terms of structure and action, and can bind to estrogen receptors in the body, showing estrogenic effects (33). In an experiment, 17-b estradiol and genistein increased the gene expression of prothrombin, factor (F) VII and fibrinogen in the treated groups compared to the control group (30, 34).

Virtual examinations of the drug effects, a well-known computational method with high accuracy, can be employed to identify the best and most stable compounds for the proposed drug. Recent developments in statistical algorithms have provided the possibility of predicting drug toxicity, absorption, and physicochemical properties relatively quickly and accurately (35, 36).

*Caenothus americanus* is a small shrub belonging to the Rhamnaceae family (37) with hemostatic properties attributed to its alkaloids (38), which interact with antithrombin and thromboplastin in the blood coagulation pathway. The exact mechanism of this process has not been precisely determined yet, but it is believed that several alkaloids in the root bark tincture trigger blood coagulation. In this framework, the decoctions of the root bark and the fresh root have been used to prevent wound bleeding (38) and can be effective in reducing clotting time (CT) (39). Also alkaloids extracted from *Zingiber officinale* and *Thymus vulgaris* have been found to reduce blood CT (40, 41). Additionally, Alkaloids cause the release of epinephrine (adrenaline) (42) and subsequently increase the amount of FV (43). Flavonoid compounds in the C. americanus could increase the expression of immune-related genes such as tumor necrosis factor (TNF)-α (39, 44–46) which in turn activates platelet aggregation and thereby reducing of clotting time (47). This gene also can activate platelets by stimulating the arachidonic acid pathway (47). Tannins present in this plant extract precipitate blood proteins (39), leading to the synthesis of thromboxane A2 and recruitment of additional platelets to form platelet aggregation at the site of damaged vessels (48). The hydroalcholic extract of the powdered root of C. americanus contains four dicarboxylic acids, all of which have blood coagulation activity (37).

In agreement with these data, C. americanus was identified as one of the effective coagulation plants. In this study, computational analyses were employed to identify the compounds of *C. americanus* with blood coagulant potential using physicochemical and pharmacokinetic servers and molecular docking techniques.

## Materials and methods

### Selection of the herbal plant and Ligand preparation

The herbal plant C. americanus was chosen for this study due to the presence of compounds with potential coagulation factor effects (Table 1). The two-dimensional (2D) structure of identified compounds in C. americanus extract was obtained from the PubChem web servers in sdf format (https://pubchem.ncbi.nlm.nih.gov/).

**Table 1.** The smiles and structure of C. americanus extract compound.

| Ligand | Smiles |
| --- | --- |
| Malic acid | O=C(O)C(=O)CC(O)O |
| Malonic acid | O=C(O)CC(=O)O |
| Oxalic acid | O=C(O)C(=O)O |
| Succinic acid | O=C(O)CCC(=O)O |

**Table 2.**
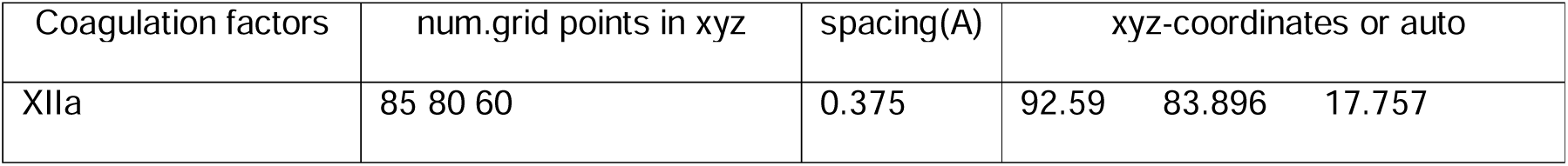
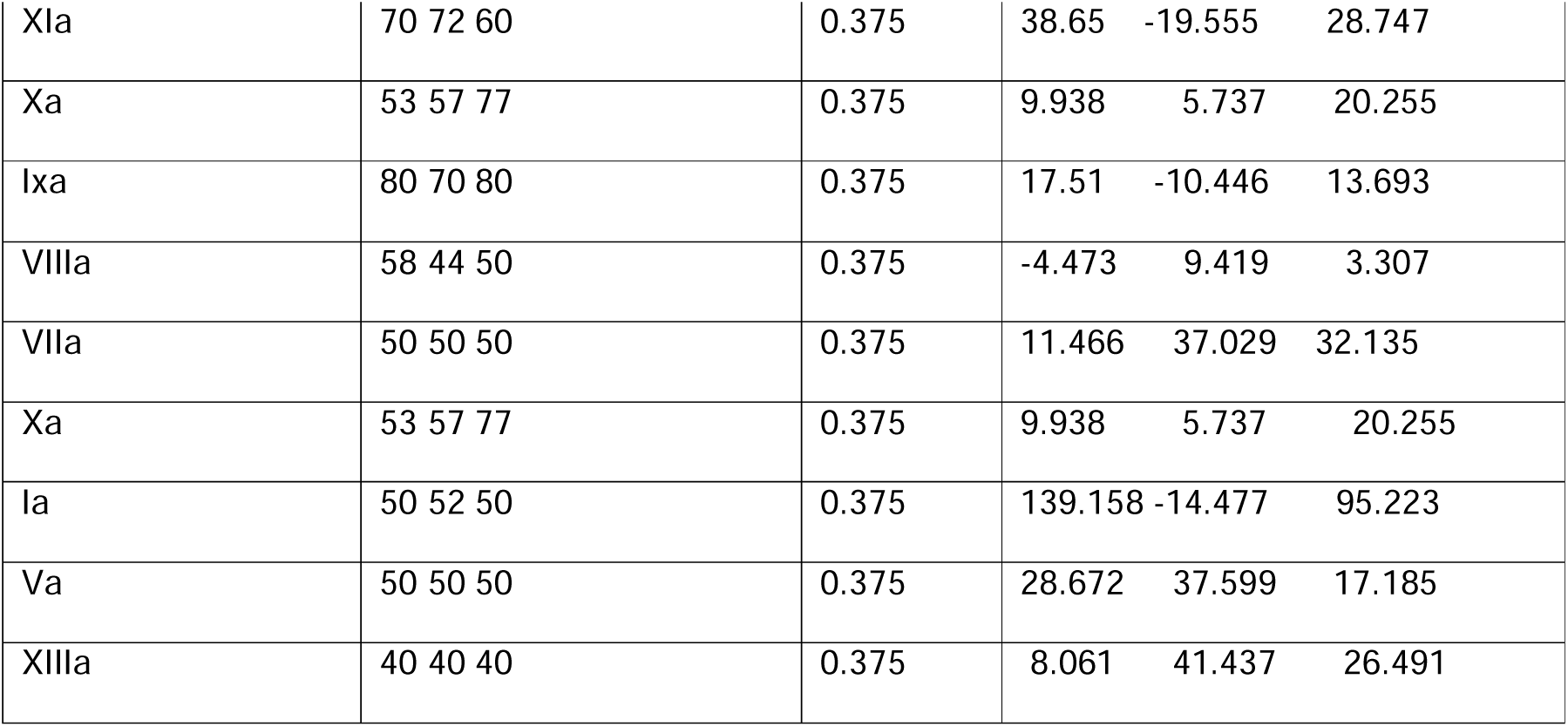
The XYZ box size used for all ligands in molecular docking.

Then, OpenBable program was used to convert the compounds to simplified molecular input line entry system (SMILES) format (Table 1). To optimize the structure of compounds and find the most stable state and the binding energy of the structures, HyperChem software was used with an Amber force field at 300K and 1000 picoseconds. The structures were prepared for AutoDock input for docking analysis after the addition of partial charge and polar hydrogens and removal of H2O.

### Pharmacokinetic properties

In the present research, four compounds of C. americanus were selected based on previous literature for the evaluation of their pharmacokinetic properties (Table 1).

The ADME-related physicochemical properties of malic acid, malonic acid, oxalic acid, and succinic acid were predicted by the Swiss ADME online web server and ADMET lab server. The SMILES format of each compound was obtained from PubChem web servers and OpenBable and then loaded to SwissADME (http://www.SwissADME.ch/) and ADMET servers (https://ADMETmesh.scbdd.com/) to calculate their physical-chemical, pharmacokinetic, and pharmacodynamics properties. The physicochemical properties, lipophilicity and solubility of the each compounds were considered for the analysis (49).

Based on gold standards and previously published research on different rules of oral bioavailability, the key parameters evaluated were MW, lipophilicity (M log P), TPSA, HBD, HBA, MR, nRot, and solubility in water, using SwissADME and ADMET lab servers. The Lipinski properties of the compounds were also investigated. According to the Lipinski rule, adequate absorption and permeability for an oral drug are likely if the molecular weight is ≤500, the oil/water distribution coefficient (LogP) ≤5, hydrogen bond donors (OH + NH count) ≤5 (expressed as the sum of OHs and NHs), and hydrogen bond acceptors (O + N atom count) ≤10 (50, 51). MW is one of the key features in small molecule drug discovery. This parameter can impact various molecular events such as absorption, bile elimination rate, blood-brain barrier (BBB) penetration, and interactions with targets (on- and off-targets) while it is also commonly monitored during the drug optimization steps (51).

### Pharmacodynamics properties

Human oral bioavailability (HOB) indicate two parameters, F20% and F30%, where positive results for both show component bioavailability in the human body up to 20%, negative results, for both show no bioavailability either 20% or 30% and positive results for F30% indicate high oral bioavailability, and negative for F20% means low oral bioavailability (52). The BOILED-Egg model was used to predict compounds’ permeability of the BBB penetration and capability of gastrointestinal (GI) absorption (49).

Plasma fraction unbound plasma (Fu) and volume of distribution (VDss) are two important parameters in drug distribution. In this respect, absorption, distribution, metabolism, excretion, and toxicity (ADMET) properties of all compounds were calculated based on the physicochemical properties determined by employing ADMETlab (ADMETLAB2.0 https://ADMETmesh.scbdd.com) and SwissADME (http://www.SwissADME.ch/).

In silico data for major human cytochrome P450 (CYP) isoforms involved in the drug metabolism, such as CYP1A2, CYP2C9, CYP2C19, and CYP3A4 were also generated to determine the excretion routes of compounds. In drug attrition, the safety profile of the compounds comes under major parameters (49) with evaluation of some safety parameters such as human hepatoxicity, carcinogenicity, oral toxicity to mice, drug-induced liver damage, and human ether-a-go-go related gene protein (HERG).

### Preparation of the target proteins

The extrinsic FVII and also factors XII, XI, IX, and VIII are considered intrinsic pathway factors. The common pathway protein factors include clotting factors V, II, XIIIa, and Xa.

The crystal structures of the coagulation factors including FIIa (PDB ID 5AFZ), FVa (PDB ID 1SSD), FVIIa (PDB ID 1W7X), FVIIIa (PDB ID 3HNY), FIXa (PDB ID 3LC3), FXa (PDB ID 2vh0), FXIa (PDB ID 5QTW), FXIIa (PDB ID 6B77), and FXIIIa (PDB ID 1F13) were obtained from the protein data bank (PDB). The initial protein structures were prepared with ArgusLab server and then optimized by removing non-bonded atoms and non-polar hydrogens using AutoDock Tools software.

### Molecular docking

Molecular docking was employed to evaluate components’ binding affinities and active binding residues estimation using AutoDock 4 software. According to the research of previous researchers and the examination of these factors in the protein bank, the active pocket of this protein was identified and the grid box was considered for the active position of each protein (Table2). The number of genetic algorithm (GA) runs for each 50 docks was selected by 25 * 106 energy measurements.

After docking, results of 50 runs were categorized into clusters and the cluster with the highest sub- rank and the binding energy level was selected for further analysis. The complex of ligand and protein was created in the selected rank and saved in pdbqt format, and Discovery Studio software was then used for visualization.

## Results

### Structural and physicochemical properties

Physicochemical properties, such as cell membrane properties and drug transport forces, are factors that affect the pharmacokinetics of drugs (53).

The reduction of risk of drug wear is related to physicochemical properties. Most of the introduced drug candidates fail at some stage of development. Over the past few years, significant growth has been observed in the understanding of the links between the physical properties of drugs and drug erosion. Evidence suggests that drug metabolism and pharmacokinetics (DMPK) and toxicity outcomes are related to the physical properties of compounds under the control of medicinal chemists and other project scientists in drug discovery projects (49, 54).

The MW of the compounds range from 90 to 134 g/mol (Table 3), and all the experimental compounds were within the optimal molecular weight range (<500).

**Table 3.** Physicochemical properties of C. americanus extract compounds.

| Ligand | Molecular Weight( g/mol) | MlogP | TPSA (Å<sup>2</sup>) | nHD | nHA | MR(c m<sup>3</sup>) | nRot | Ali log S |
| --- | --- | --- | --- | --- | --- | --- | --- | --- |
| Malic acid | 134.020 | -1.37 | 94.830 | 3 | 5 | 26.05 | 3 | -0.24 |
| Malonic acid | 104.010 | -0.99 | 74.600 | 2 | 4 | 20.08 | 2 | -0.28 |
| Oxalic acid | 90 | -1.51 | 74.600 | 2 | 4 | 15.27 | 1 | -0.86 |
| Succinic acid | 118.030 | -0.54 | 74.600 | 2 | 4 | 24.89 | 3 | -0.51 |
| Molecular weight (MW) <500 (g/mol), Moriguchi octanol-water partition coefficient (MlogP) <5, topological polar surface area (TPSA) ≤140 Å<sup>2</sup>; number of hydrogen bond donors (nHD,) <5, number of hydrogen bond acceptors (nHA) <10; Molar Refractivity (MR) (optimal: 40 and 130), number of rotatable bonds (nRot) (optimal: 0~11); aqueous solubility descriptor (Ali log S) ≤ 0 |  |  |  |  |  |  |  |  |

Molecular weight (MW) <500 (g/mol), Moriguchi octanol-water partition coefficient (MlogP) <5, topological polar surface area (TPSA) ≤140 Å<sup>2</sup>; number of hydrogen bond donors (nHD,) <5, number of hydrogen bond acceptors (nHA) <10; Molar Refractivity (MR) (optimal: 40 and 130), number of rotatable bonds (nRot) (optimal: 0~11); aqueous solubility descriptor (Ali log S) ≤ 0

Log p expresses the lipophilicity of the drug and plays an essential role in determining several ADMET parameters, as well as potency. Lipophilicity which was calculated in this study with Mlog p values has an important role in determining several parameters of ADMET as well as potency. Permeability decreases at very low amounts of lipophilicity, while solubility and metabolism are more compromised in high amounts of lipophilicity. Very hydrophilic compounds may penetrate into membranes by being attached to the bilayer membrane, while highly lipophilic compounds are usually unable to passively diffuse through membranes because they hardly enter the hydrophobic interior of the lipophilic bilayer. Toxicity-related issues such as the ability to hERG inhibition, phospholipidosis, or ability to CYP inhibitions are problematic for compounds with high lipophilicity (51). The log P value of compounds ranged from -1.51 to -0.54 (Table 3). Therefore, these compounds can cross the membrane by simple diffusion (53).

Polar surface area (PSA) is a useful molecular descriptor in the study of drug transport characteristics in the body, such as intestinal absorption and penetration through the BBB. The measurement of PSA is complicated by the need to accurately calculate of the appropriate 3-dimensional molecular geometry or set of geometries for each drug under study (55). The topological PSA (TPSA) of all compounds ranged from 73 to 94. According to the cutoff values set for TPSA (≤140 Å^2^) by Veber et al (56), 95% of our compounds were to have a high probability of oral bioavailability.

Hydrogen bond acceptors and donors are other important factors related to drug polarity and permeability (51). Our analysis revealed that the number of HBDs in compounds ranged from 2 to 3, while the number of HBAs ranged from 4 to 5. All the compounds in this study satisfied Lipinski’s rule, HBA < 10.

One of the important parameters for the oral bioavailability of drugs is MR described by the Ghoses rule (57). Concordantly, Lipinski’s rule states that the value of MR should lie between 40 and 130 (cm^3^) for drug-likeness (49). Among the studies of compounds, no compound had been included in Ghose’s law.

Moreever, calculating the number of links around which rotation is possible is one of the important parameters in Veber’s rule (58). Our analysis showed that the number of rotatable bonds of all compounds was within the optimal range (Table 3).

Solubility in the intestinal fluid is also another important parameter of oral drugs because decreased solubility may limit intestinal absorption through the portal vein system (51). Solubility (log s) of our compounds was in the optimal range (Table 3).

### Drug-likeness rules based on physicochemical properties

In the early stages of drug discovery or while preparing a library of suitable chemical compounds for drug discovery, certain principles were established for selecting compounds. Among the first applications of personal computers in the drug discovery, Lepinski et al. in 1997 formulated the Rule of 5 (51).

Since all of the compounds displayed characteristics in agreement with Lipinski’s properties, they are expected to have no problems in the absorption and permeability process and thus this compounds can be subjected for further preclinical trials (Table 3).

### Permeability and absorption of compounds

Caco2 (human colorectal epithelial adenocarcinoma cell lines) is mainly used as laboratory models for predicting and calculating the absorption of an oral drug in the human intestinal mucosa (58). The results from ADMETlab and SwissADME web servers show that none of compounds have the ability to pass through monolayers of Caco-2 cells (Table 4).

**Table 4.** Properties of absorption, distribution and excretion of C.americanus extract compounds. Some properties related to absorption, distribution and excretion; **Absorption**, Caco-2 Permeability: Optimal-higher than - 5.15 Log units (a); MDCK permeability, low permeability: < 2 × 10−6 cm/s, medium permeability: 2–20 × 10−6 cm/s, high passive permeability: > 20 × 10−6 cm/s (b) (60); Bioavailibity: F20% + (bioavailability < 20%), Category 0: F20%- (bioavailability ≥ 20%); F30%: 30% Bioavailability (Category 1: F30% + (bioavailability< 30%); Category 0: F30%- (bioavailability ≥ 30%)[50]**; Distribution,** Plasma protein binding: Plasma protein binding classification computed by ADMETlab server. PPB, Plasma Protein Binding (Optimal: > 90%)(a); VD, Volume Distribution (VD<0.07l/Kg correspond to bind with plasma protein or highly hydrophilic, VD: 0.07-0.7 L/Kg corresponds to evenly distributed and VD > 0.7 L/Kg corresponds to distribution towards tissue components)(b); Fu, the fraction unbound in plasma (Low: <5%; Middle: 5∼20%; High: > 20% (c) (60, 61); **Excretion**, Half-life and clearance rate CL: Clearance (High: >15 mL/min/kg; moderate: 5-15 mL/min/kg; low: <5 mL/min/kg) (a); T1/2, Half-life (Category 1: long half-life; Category 0: short half-life; long half-life: >3h; short half-life: <3h) (b).(60); Mal, malic acid; Malo, malonic acid; Oxa, Oxalic acid; Suc, Succinic acid.

| Ligand | absorption |  |  |  | Distribution |  |  | Excretion |  |
| --- | --- | --- | --- | --- | --- | --- | --- | --- | --- |
|  | caco cells | MDCK(cm/s) | Bioavailability |  | PPB(%) | VD (L/Kg) | FU(%) | CL(mL/min/kg) | T1/2(Hourse) |
|  |  |  | F20% | F30% |  |  |  |  |  |
| Mal | -5.967 | 0.0075 | + | + | 16.822 | 0.278 | 76.029 | 3.028 | 0.852 |
| Malo | -5.858 | 0.0028 | - | - | 18.209 | 0.244 | 65.691 | 4.386 | 0.849 |
| Oxa | -5.896 | 0.0017 | + | + | 15.303 | 0.292 | 67.469 | 1.754 | 0.893 |
| Suc | -5.903 | 0.0029 | - | - | 31.728 | 0.253 | 57.44 | 4.976 | 0.842 |

MDCK cells are also among the models used for predicting drug permeability in humans. These cells are advantageous over Caco-2 cells as they can form a single layer with strong intracellular connections in three days and are considered a suitable screening system to check intracellular passage (59). Our analyses in ADMET and SwissADME server showed that all the compounds have high absorption in the MDCK cells (Table 4).

### Bioavailability

Human oral bioavailability (HOB) plays a significant role in determining the success of new drugs in clinical trials. However, measuring HOB using experimental tests is expensive and time-consuming. Therefore, the use of computational models to evaluate HOB before synthesizing new drugs could help in finding effective medicine. Fig 1 displays Bioavailability Radars, which provide better visualization of the effect of geometrical and structural properties on the bioavailability of the compounds in Table 3.

**Figure 1.**
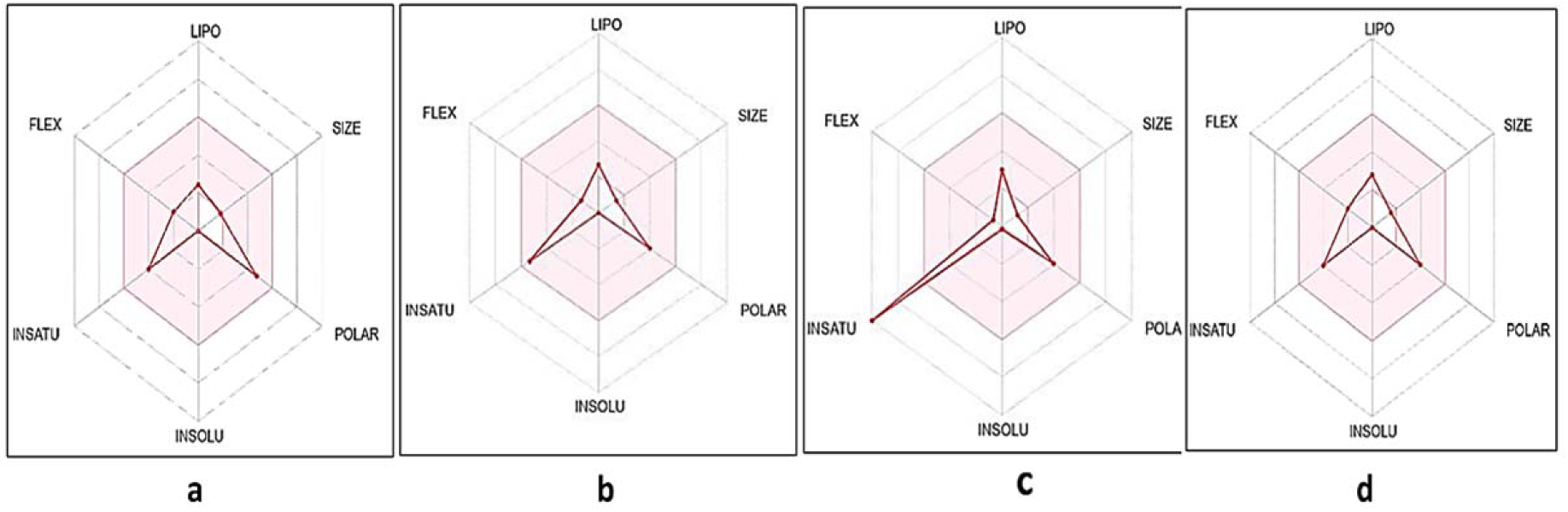
Bioavailability radar (pink area exhibits an optimal range of particular property) for the studied compounds. [LIPO = lipophilicity as XLOGP3; SIZE = size as molecular weight; POLAR= polarity as TPSA (topological polar surface area); INSOLU = insolubility in water by log S scale; INSATU = insaturation as per fraction of carbons in the sp3 hybridization and FLEX = flexibility as per rotatable bonds] (63). The colored area (pink) is the most favorable part for each of the bioavailability features (LIPO, SIZE, INSOLU, POLAR, INSATU, and FLEX) of the drug (62). a, malic acid; b, malonic acid; c, oxalic acid; d, succinic acid.

Malic acid and oxalic acid compounds are bioavailable in the human body up to 20% and 30%, respectively (Table 4). The oral bioavailability of the compounds in plants and chemical drugs was evaluated using the SwissADME web tool, and the results are presented in Fig 1. The LIPO (Lipophilicity) of the compounds was determined using the XLOGP value. Surprisingly, all the compounds fell within the colored region and the LIPO recommended range of -1.26 to -0.25. According to the Lipinski rule, a good drug should have a molecular weight of less than 500 g/mol (62). The INSOLU (insolubility) requirement of the compounds as depicted in their ESOL (LogS) and ESOL Class revealed that all compounds was very soluble (62).

The Total Polarity Surface Area (TPSA) with an optimal range of 20 and 130 A° was used to calculate the POLAR of the compounds (62). As shown in Fig 2, all the compounds have a considerable amount of polarities. To determine the unsaturation (INSATU) and flexibility (FLEX) of compounds, rotatable bonds must be less than 9. Interestingly, all the compounds fell within the FLEX recommended range of values. Put together, malic acid (a), malonic acid (b), and succinic acid (d) had the best oral bioavailability, since all their physicochemical properties fell within the optimal colored region (pink).

**Figure 2.**
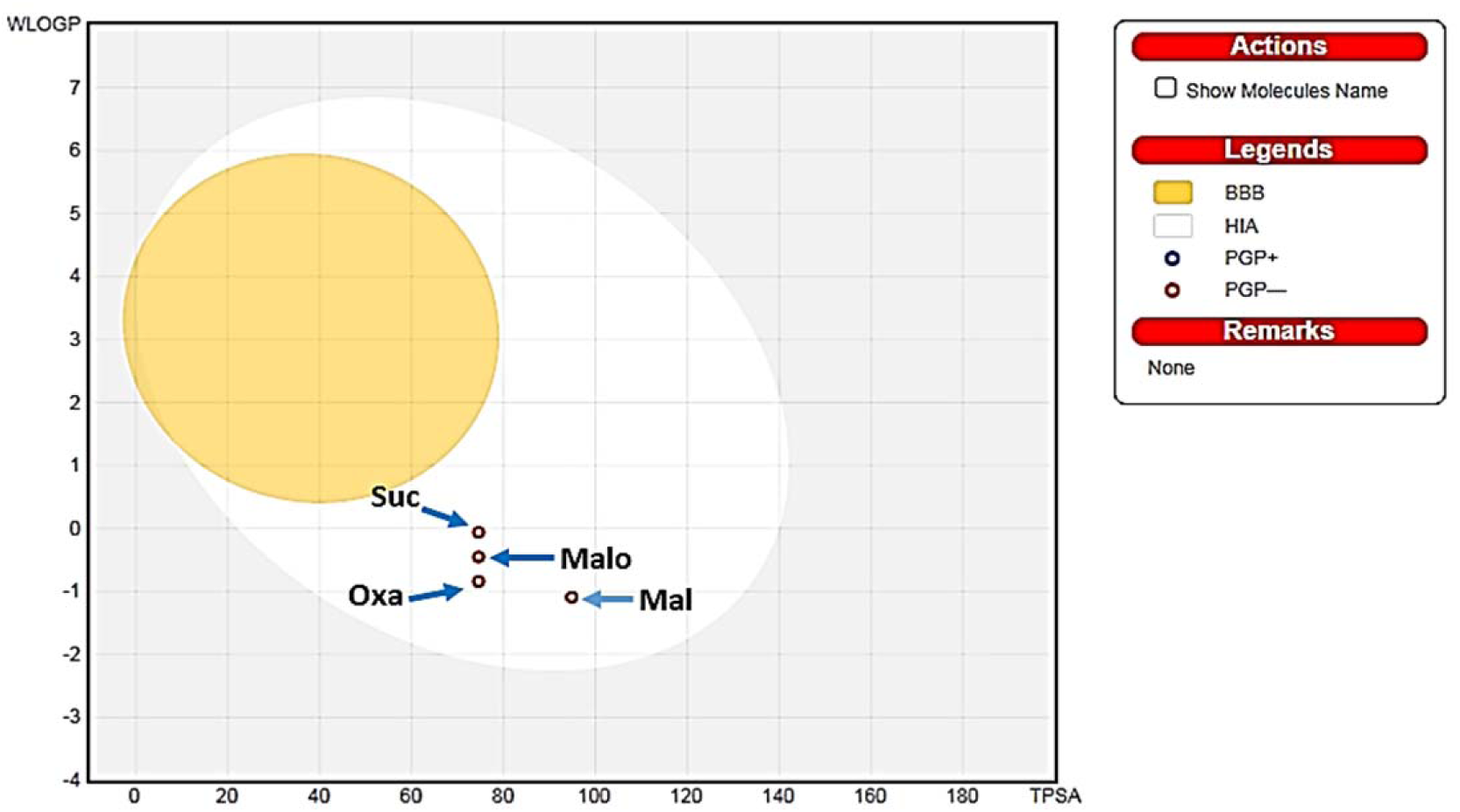
BOILED-Egg ADMET model all compounds. Gastrointestinal absorption or human intestinal (HIA); blood-brain barrier (BBB) penetration; (PGP) substrate for P-glycoprotein; (PGP-) Not a substrate for P-glycoprotein. Mal, malic acid; Malo, malonic acid; Oxa, Oxalic acid; Suc, Succinic acid.

### Distribution results

Drug distribution includes the distribution of the compound into various parts of the body (64). In this study, various parameters that can be accurately investigated in silico, such as the ability to bind to blood plasma proteins, the volume of distribution, the unbound fraction in the plasma, and the ability to cross the BBB were taken into account.

#### Plasma Protein Binding (PPB)

The PPB parameter refers to the binding of a drug to plasma proteins and lipids after it has been distributed in circulating blood. The free drug theory that only the free drug can enter tissues, reach the target sites, and exert its effect for a useful period of time (65).

According to the results of in silico evaluation using ADMETlab, all compounds had poor ability to bind to plasma proteins (Table 4).

#### Volume of Distribution (VD)

The steady-state volume of distribution (VDss) is a crucial pharmacokinetic parameter that describes the distribution of an injected amount of drug to create a concentration equivalent to that in blood plasma (58). In other hand, the VD is the volume of plasma or blood in which the compound appears to be dissolved at a steady state or equilibrium (60, 66).

A VD in L/Kg indicates whether a sufficient drug concentration can be maintained in the bloodstream and can be calculated from the distribution of a drug in circulation relative to the distribution in the rest of the body. A VD <0.07 L/kg indicates high binding to plasma proteins (observed for highly hydrophilic compounds), a value between 0.07-0.7 L/kg corresponds to an even distribution between the two compartments, and a VD >0.7 L/kg corresponds to distribution towards tissue components (observed for highly lipophilic compounds) (50). Our analysis indicated that all compounds showed an even distribution between compartments (Table 4).

#### Fraction unbound plasma (FU)

Predicting the FU provides a good understanding of the pharmacokinetic features of a drug, which can help with candidate selection in the early stages of drug discovery. This parameter is a key determinant of drug efficacy in pharmacokinetic and pharmacodynamics studies. In general, the unbound (free) drug can interact with pharmacological target proteins such as channels, receptors, and enzymes and is able to diffuse between plasma and tissues (67). Based on the parameters of ADMETlab and SwissADME, all compounds had a high FU (Table 4). According to the results, an equilibrium relationship between PPB and FU was found, where a decrease in the binding of the drug to plasma proteins led to an increase in its unbound fraction and vice versa.

#### BOILED-Egg for Prediction of GI Absorption and BBB Permeability

GI absorption and BBB Permeability are crucial factors in drug development. The BOILED-Egg model can accurately calculate the polarity and lipophilicity of compounds and provide data with speed, and clear graphic outputs.

This model predicts a high probability of passive absorption of compounds by the GI tract, which is represented by the white region, whereas the BBB permeability is displayed in yellow. The red color indicator shows the nonsubstrate of Pgp, represented as (PGP−), whereas, the blue color represents molecules actively effluxed by p-glycoprotein (PGP+).

Based on GI incidence, which predicts drug absorption in the intestine, all compounds in this study were predicted to be absorbed into the intestine (Fig 2). However, drug penetration into the BBB is an unwanted side effect of drugs that are not intended to target the central nervous system (60).

Predictive assessment using ADMETlab and SwissADME indicated that none compounds can cross the BBB penetration (Fig 2).

P-glycoprotein (P-gp) is a member of the ATP-binding cassette (ABC) transmembrane superfamily (58). It is one of the most important parameters of ADMET, and plays a vital role in protecting cells against toxic material such as drugs and toxins by reducing the absorption of xenobiotic and extruding these compounds (66).

P-gp is preferentially expressed in important body tissues such as brain endothelial capillaries containing the BBB to investigate distribution. Molecular weight (MW)>400 and log P>4 are preferentially transported by P-gp (68). Our analyses indicated that none of the compounds were substrates of p-glycoproteins (Fig 2). Considering that the compounds in Table 3 had a molecular weight of less than 400 and log P of less than 4, they were not glycoproteins’ substrates, and their transfer is not done by glycoproteins, on the other hand, they cannot cross the BBB through P-gp.

### Metabolic properties

The metabolic process is capable of metabolizing the drugs, which require processing by redox enzymes, including the cytochrome P450 system (CYPs) (60). Drug metabolization by CYPs is essential for detoxification to prevent the drugs accumulation in the body. In this respect, the p450 cytochrome plays an important role in the study and investigation of compound metabolisms, and its subunits including CYP1A2, CYP2C9, CYP2C19, and CYP3A4 are responsible for metabolizing more than 90% of drugs (58). Inhibition of these cytochromes is one of the major causes of interactions related to drug pharmacokinetics, which leads to a variety of side effects, including drug toxicity due to lower clearance and drug accumulation (57). Approximately 8.9% of drugs subject to biotransformation in the liver are metabolize under the action of the CYP1A2 enzyme (58). Prediction of drug decomposition using ADMETlab and SwissADME indicated none of compounds were potent inhibitors of CYP1A2 (Table 4).

Approximately 12% and 6.8% of drugs subject to biotransformation in the liver are metabolized Under the action of the CYP2C9 and CYP2C19 enzyme, respectively (58). According to the prediction results in ADMETlab and SwissADME, none of the compounds inhibited these cytochromes (58). Approximately 20% and 30.2% of drugs subject to biotransformation in the liver are metabolized Under the action of the CYPD6 and CYP3A4 enzymes, respectively (58). The online servers ADMETlab and SwissADME predicted that also none of the compounds were potent inhibitors of these cytochromes (Table 5).

**Table 5.** Prediction of toxicity and methabolic. hERG Blockers: Category 1: active, Category 0: inactive; H-HT: Human Hepatotoxicity (Category 1: H-HT positive(+); Category 0: H-HT negative(-)); DILI: Drug Induced Liver Injury (Category 1: drugs with a high risk of DILI; Category 0: drugs with no risk of DILI); AMES Toxicity: Category 1: Ames positive(+), Category 0: Ames negative(-); Rat Oral Toxicity: Category 0: low-toxicity, Category 1: high-toxicity; Respiratory Toxicity: Category 1: respiratory toxicants, Category 0:respiratory nontoxicants; Skin Sensitization: Category 1: Sensitizer, Category 0: Nonsensitizer (60).

| Ligands | Toxicity |  |  |  |  |  |  | Metabolic |  |  |  |  |
| --- | --- | --- | --- | --- | --- | --- | --- | --- | --- | --- | --- | --- |
|  | hERG Blockers | H-HT | DILI | AMES Toxicity | Rat Oral Acute Toxicity | Respiratory Toxicity | skin sensitization | CYP inhibitor |  |  |  |  |
|  |  |  |  |  |  |  |  | 1A2 | 2C19 | 2C9 | 2D6 | 3A4 |
| Malic acid | - | - | - | - | - | - | - | - | - | - | - | - |
| Malonic acid | - | - | + | - | - | + | + | - | - | - | - | - |
| Oxalic acid | - | - | + | - | - | - | + | - | - | - | - | - |
| Succinic acid | - | - | - | - | - | - | - | - | - | - | - | - |

### Excretion (half-life and clearance)

The excretion of drugs/molecules is predicted based on two parameters, half-life (t1/2) and clearance (CL), on the ADMETlab server (60). The t1⁄2 is a parameter that represents the time required for the plasma concentration of a drug to reduce to half of its initial value (58). According to the predictions, the compounds of this study had a low half-life. CL is a drug parameter that expresses the rate of irreversible drug removal from the body (58). The online server predicted that the CL value of all compounds was < 5 mL/min/kg (Table 4).

### Toxicity

Developing a new drug without toxic properties is crucial. Hence, the potential toxicity of C. americanus compounds has been evaluated using ADMETlab, with eleven computational toxicity parameters, in various animal and human organs (66, 69). Human ether-a-go-go related gene (HERG) is involved in the fatal arrhythmia known as torsade de pointes or the long QT syndrome, encoding a potassium ion (K+) channel (70). The HERG K+ channel participates in the electrical activity of the heart, which coordinates cardiac warming. interestingly, investigating the effect of toxicity of compounds on the electrical activity of the heart has therapeutic value (70). According to the prediction results from the ADMETlab online server, none of the compounds inhibited the potassium channels of the heart (Table 5). Hepatotoxicity is one of the leading causes of drug withdrawal from pharmaceutical development and clinical use (64). The major reason for drug non-approval by the US food and drug administration is drug-induced hepatotoxicity (71). According to the prediction results from the ADMETlab online server, none of the compounds caused human hepatotoxicity (Table 5). Drug-induced liver injury accounts for 10% of all cases of acute hepatitis, 5% of all hospitalization, and 5% of all acute liver failure (71). In predicting DILI parameter with the ADMET server, all compounds (except malonic) did not cause liver damage (Table5). The Ames test is used to determine the mutagenic effects of drugs (72). According to the prediction results from the ADMETlab online server, none of the compounds were mutagenic (Table 5).

The median lethal dose parameter, LD50, is used to measure acute toxicity in mice, which determines the relative toxicity of a compounds (66). The prediction of drug decomposition using ADMETlab indicated none of the compounds caused oral toxicity in rats. Additionally, the prediction of drug decomposition using ADMETlab indicated none of the compounds (except malonic acid) were harmful to the respiratory system. According to predictions, malic acid and succinic acid were the only compounds that will not cause skin sensitivity (Table 5).

### Molecular docking analysis of drugs with coagulation factors of the intrinsic pathway

#### Coagulation FXIIa

Among the tested compounds, Malic acid showed the best docking scores (The binding energy -4.37 (kcal/mol)). It was observed to interact with the receptor through one (1) conventional hydrogen bond with SER-217 at a distance of 1.86Å and one (1) van der Waals bond with TRP-215 at an active site of the receptor (see Fig 3, and supplementary 1). Malonic acid had promising docking scores (the binding energy -4.25 (kcal/mol). This compound was found to interact with the receptor through one (1) van der Waals bond with TRP-215 (see Fig 3 and supplementary 1). Oxalic acid (binding energy –3.52 kcal/mol) was also bound to the receptor via one (1) van der Waals bond with TRP-215 (see Fig 3 and supplementary 1).

**Figure 3.**
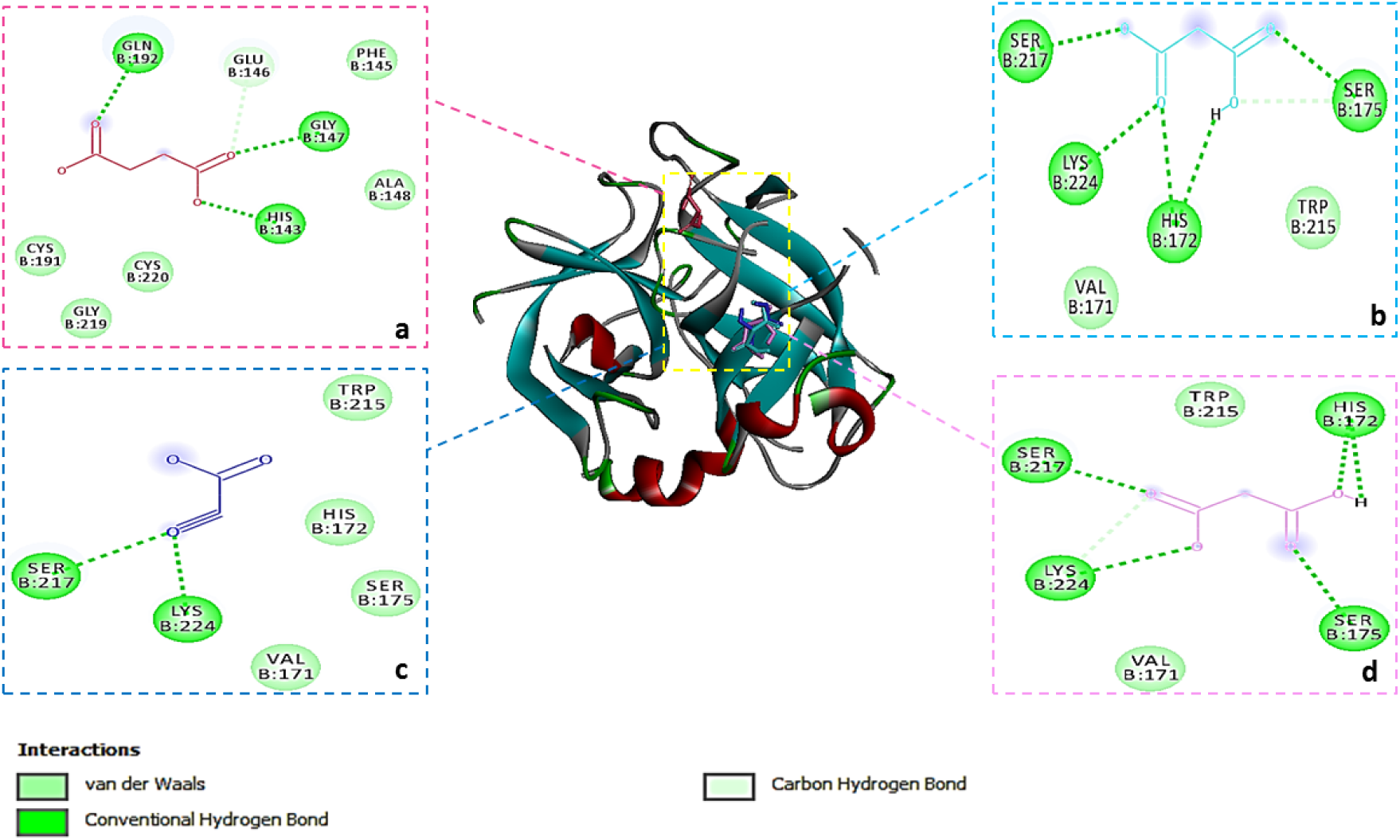
3-D (three dimensional) diagram and 2-D (two dimensional) diagram showing several interactions between succinic acid (a), malonic acid (b), oxalic acid (c), malic acid (d) and coagulation factor XIIa

The 3D and 2D representations of succinic acid (binding energy –3.92 (kcal/mol), supplementary 1) in the binding active pocket of the coagulation FXIIa receptor are shown in Fig 3. It is observed to interact with the protein receptor via one (1) conventional hydrogen bond with GLN-192 at a distance of 2.14Å and one (1) van der Waals bond with TRP-215 (see Fig 3 and supplementary 1).

#### Coagulation FXIa

The molecular docked pose of the minimum binding affinity conformer of all compounds demonstrated that they bind to the active site (73, 74) of the coagulation FXIa. Fig 4 shows that malic acid, malonic acid, succinic acid, and oxalic acid bind through conventional H-bond with LYS-192, in the S1 pocket with the binding energy of -3.83, -3.87, -3.9, and -3.74 (kcal/mol), respectively (supplementary 1). Also, these compounds had a van der Waals bond with the TYR-143 residue in the S2 pocket (Fig 4 and supplementary 1).

**Figure 4.**
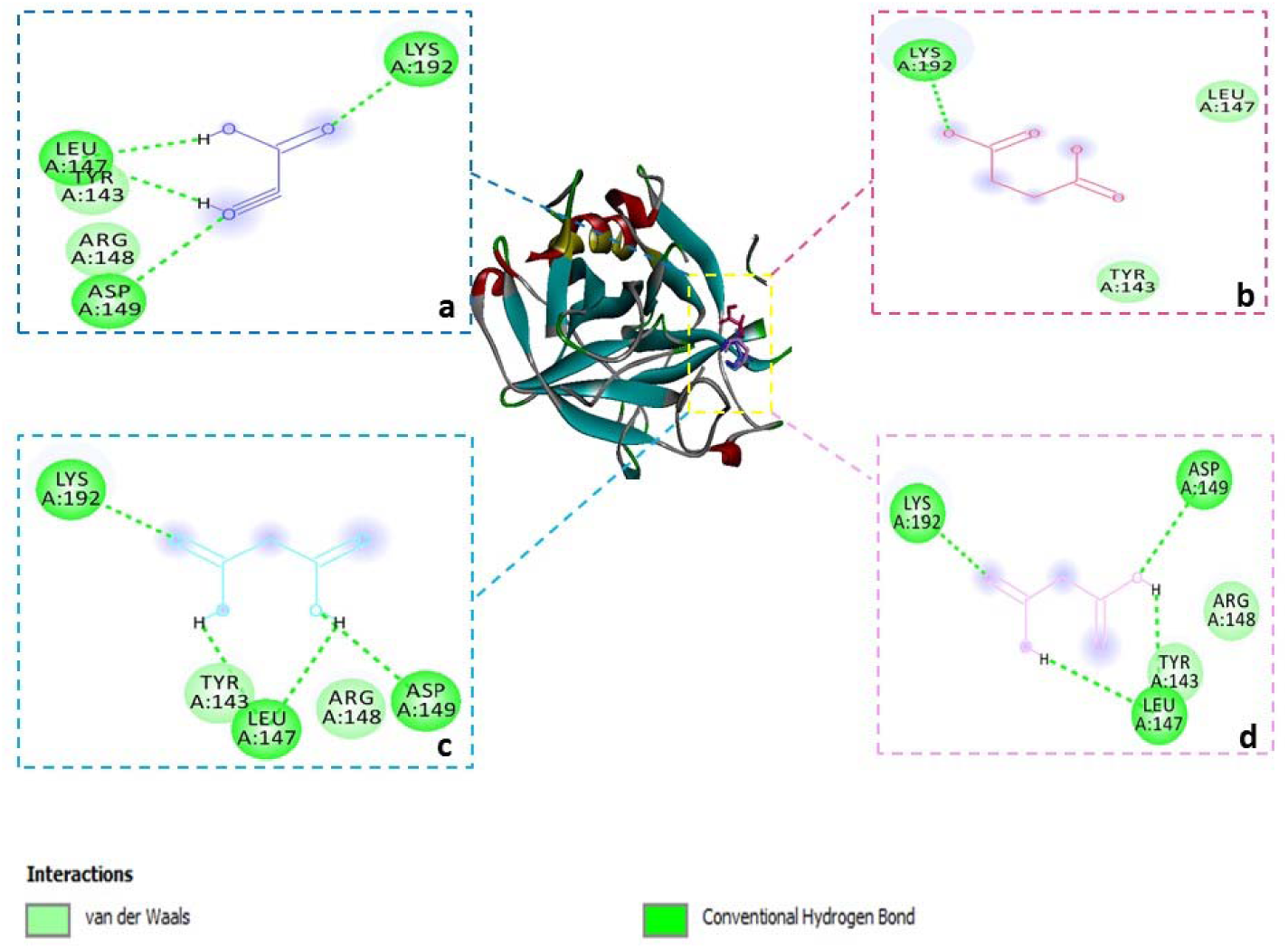
3-D (three dimensional) diagram and 2-D (two dimensional) diagram showing several interactions between oxalic acid (a), succinic (b), malonic acid (c) and malic acid (d) and coagulation factor XIa

In this study, succinic acid, one of the dicarboxylic acids in blood coagulation factor, showed a higher affinity for FXIa than the other three compounds. This compound forms both hydrogen bond and van der Waals bond at the binding site. The other three compounds also formed more hydrogen bonds in the binding site of FXIa, but their binding to FXIa was always different. Oxalic acid bound to FXIa with a higher inhibition constant (1.81 millimolar) and lower energy (-3.74 kcal/mol) than malic acid (binding energy -3.83 kcal/mol and inhibition constant 1.55 millimolar) and malonic acid (binding energy -3.87 kcal/mol and inhibition constant 1.47 millimolar). But in any case, the results of molecular docking showed that all four compounds had a high tendency to bind to amino acids of the active site through hydrogen and van der Waals bonds (see Fig 4 and supplementary 1).

#### Coagulation FIXa

The dockings of the oxalic acid showed interactions between the oxygen group and the key residues GLU-60, GLY-61, and VAL-62 of the active site in the S1 pocket (Fig 5) with an energy of -5.15 (kcal/mol) (supplementary 1). This compound formed a conventional hydrogen bond with GLU-60 residue of 2.93Å, a van der Waals bond with GLY61, and two conventional hydrogen bonds with VAL-62 residue of 1.82Å and 1.99Å in the active site of this protein, while the other three compounds (malic acid, malonic acid, and succinic acid) were placed in the serine protease position of the protease with energies of -5.57, -5.66, and -5.99 (kcal/mol), respectively. Additionally, oxalic acid with the highest inhibitory constant (168.46 millimolar) compared to the other compounds caused the release of energy 5.15 kcal/mol, which seems to be effective in protein activation (see Fig 5 and supplementary 1).

**Figure 5.**
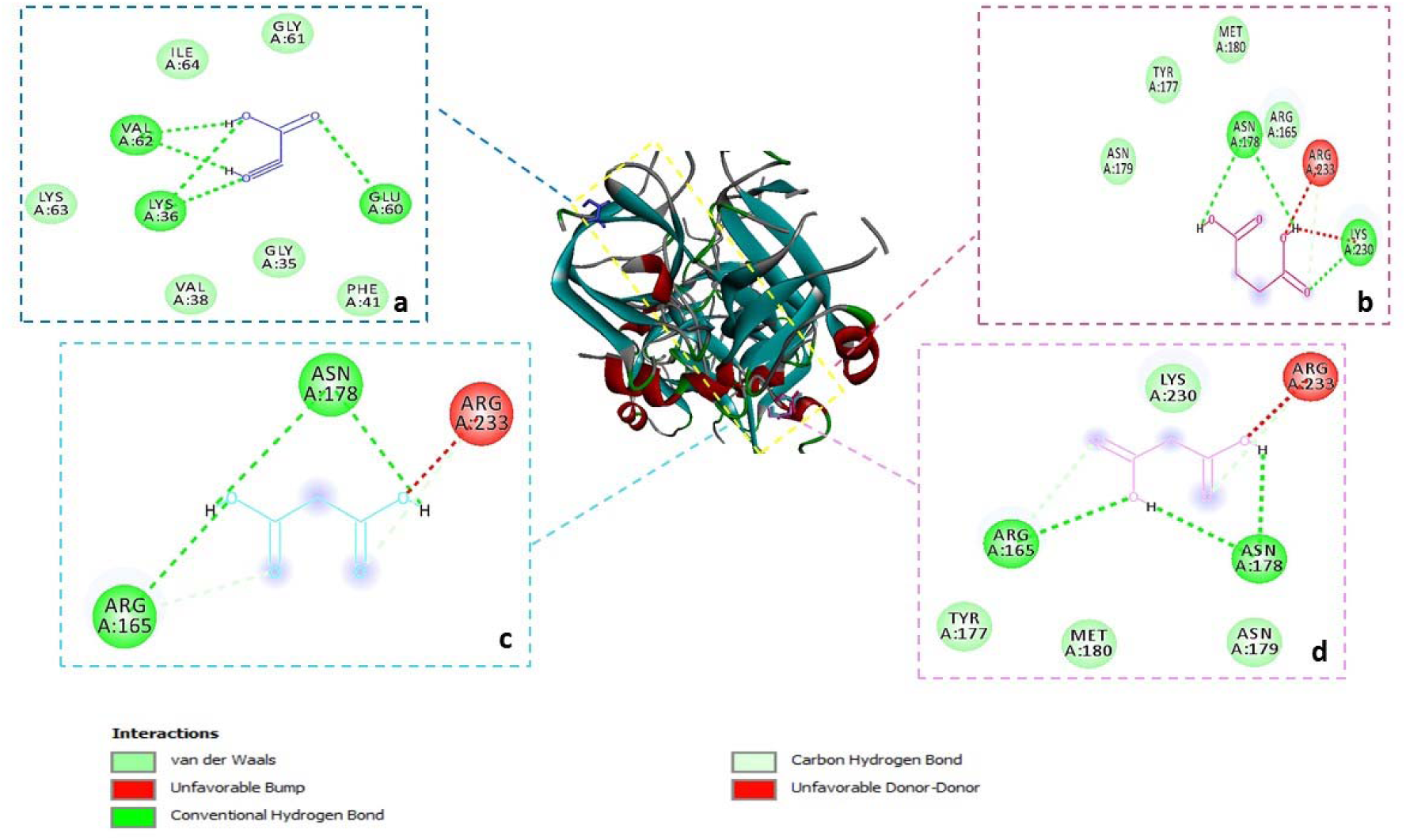
3-D (three dimensional) diagram and 2-D (two dimensional) diagram showing several interactions between oxalic acid (a), succinic (b), malonic acid (c) and malic acid (d) and coagulation factor IXa

#### Coagulation FVIIIa

FVIII is a 2332-amino-acid protein with the several domains including A1-A2-B-A3-C1-C2. Its heavy chain includes the A1 subunit (1–372), the A2 subunit (373–740), and a light chain containing the A3, C1, and C2 domains (1690–2332) (75, 76). Malic acid, malonic acid, and succinic acid interacted with the active site residues of FVIIIa in the C-domain site .

Malic acid (binding energy -2.14 kcal/mol) was linked to the active site of the receptor via one conventional hydrogen bond with PRO-2300 at a distance of 2.07Å, and two van der Waals bonds with PRO-2299 and LEU-2302 (see Fig 6 and supplementary 1). The malonic acid was the second-best ligand (energy binding -2.11 kcal/mol). This compound was linked to active site of the protein through a conventional hydrogen bond with residue PRO-300 at a distances of 2.16Å, and two van der Waals bonds with PRO-2299 and LEU 2302 (see Fig 6 and supplementary 1).

**Figure 6.**
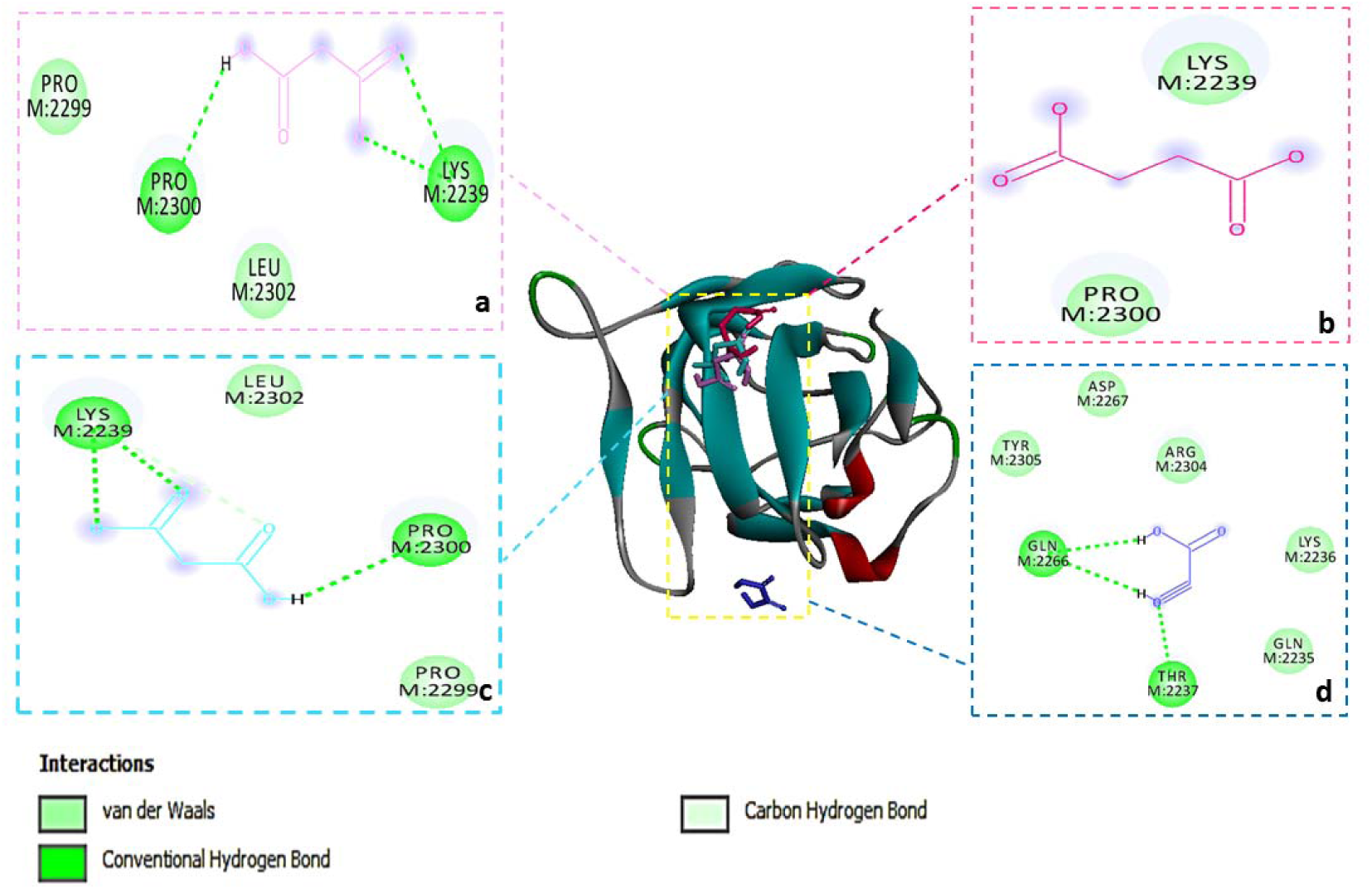
3-D (three dimensional) diagram and 2-D (two dimensional) diagram showing several interactions between malic acid (a), succinic (b), malonic acid (c) and oxalic acid (d) and coagulation factor VIIIa.

The 3-D and 2-D representations of the succinic acid (binding energy -1.88 kcal/mol) in the binding active pocket of the coagulation FVIIIa receptor are shown in Fig 6. It is observed to interact with the protein receptor via van der Waals bonds with PRO-300 (supplementary 1).

Also, the oxalic acid interacted with the binding energy of -2.52 kcal/mol near the active site in the position of domain C residues (see Fig 6 and supplementary 1).

### Molecular docking analysis of drugs with coagulation factors of the extrinsic pathway

#### Coagulation FVIIa

Four ligands, namely malic acid, malonic acid, succinic acid, and oxalic acid interacted with the residues of the S1 pocket in the active site with binding energies of -4.34, -4.35, -4.02, and -3.55 (kcal/mol), respectively. Among these, malonic acid had the best docking scores, with binding energy of -4.35 kcal/mol. It was found to bind to the receptor through two conventional hydrogen bonds with LYS-192 at a distance of 1.59Å and 1.59Å, and a van der Waals bond with CYS-191 at an active site of the receptor (see Fig 7 and supplementary 2).

**Figure 7.**
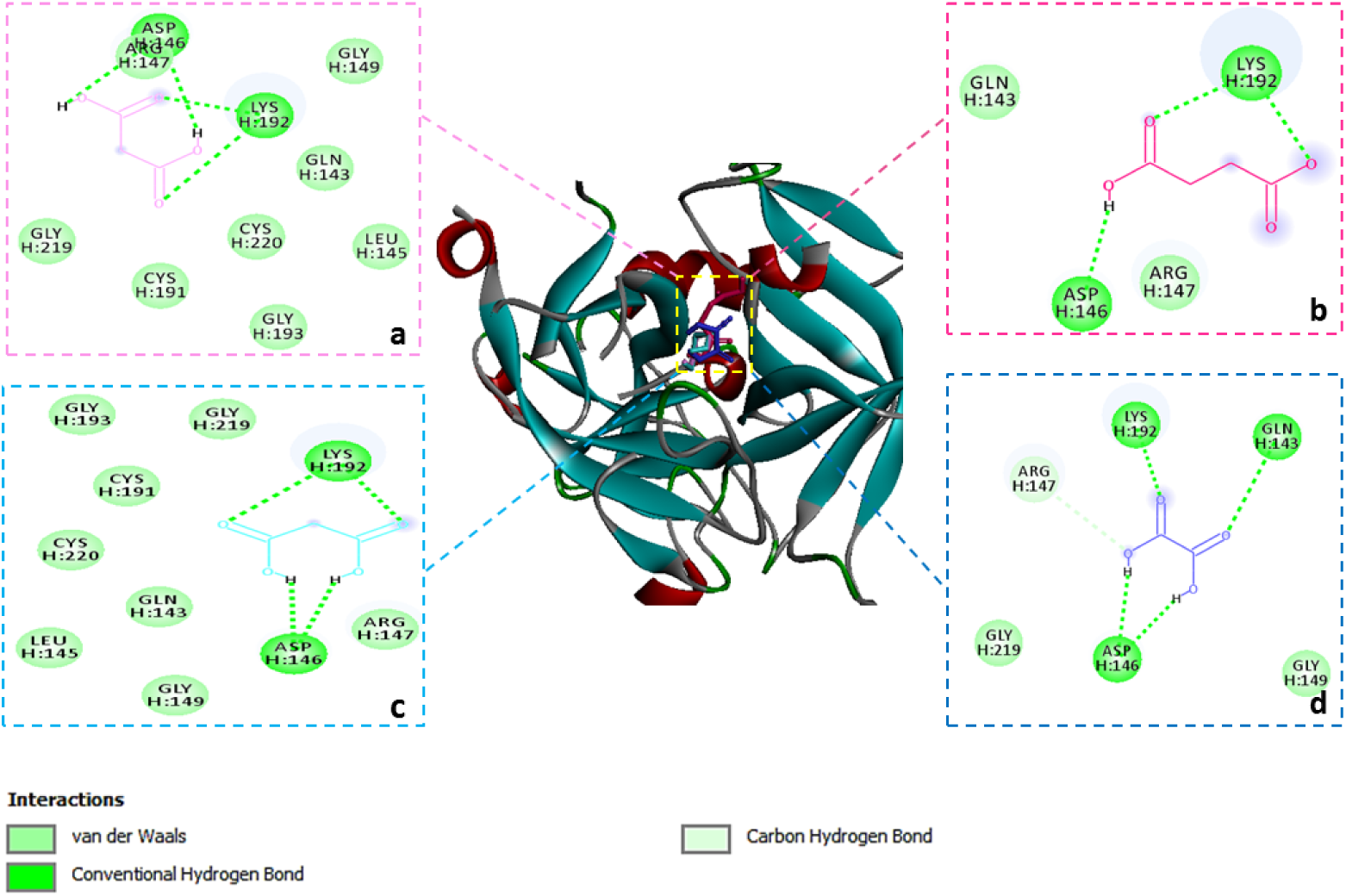
3-D (three dimensional) diagram and 2-D (two dimensional) diagram showing several interactions between malic acid (a), malonic acid (b), succinic acid (c) and oxalic acid (d) with coagulation factor VIIa

Succinic acid and oxalic acid also revealed promising docking scores with binding energy of -4.02 kcal/mol. Succinic acid was found to interact with the receptor through two conventional Hydrogen bonds with LYS-192 at a distance of 1.64Å and 1.68Å, while oxalic acid was interacted to the receptor via one conventional hydrogen bond, with LYS 192 at a distance of 1.80Å (see Fig 7 and supplementary 2). In addition to malic acid, the malonic acid, and oxalic acid interacted with S3 pocket residues in the active site (see Fig 7 and supplementary 2).

### Molecular docking analysis of drugs with coagulation factors of the common pathway

#### Coagulation FXa

The docking energy and binding residues with coagulation FXa active site are presented in supplementary 3. The heavy chain of coagulation FXa contains a serine protease domain in a trypsin- like closed â-barrel fold encompassing the active triad of Ser195-His57-Asp102 and two essential protein subsites, S1 and S4, which are frequently explored in structure-guided drug discovery (77). In the docking of FXa (Fig. 8), it was observed that four marker compounds could insert into the active pocket of FXa via various interactions, such as hydrogen bond and van der Waals, etc. The main part of malonic acid is located in the active triad of His-57 with a conventional bond length of 1.57 Å and pocket S1 (two conventional Hydrogen bonds of length 1.76 Å and 2.05Å with Ser195 residues and one conventional Hydrogen bond of length 1.91Å with Asp-194 (with a binding energy of -3.37 kcal/mol) (see Fig 8 and supplementary 3). Malic acid, succinic acid, and oxalic acid were mainly located at the S1 pocket (interaction with residue LYS-96). Malic acid and succinic acid were found to interact with the receptor via one conventional hydrogen bond with LYS-96 at distances of 1.83Å and 1.78Å respectively. Also, the oxalic acid interacted with LYS-96 with two conventional hydrogen bonds of 1.63Å and 2.08Å, respectively. The binding energy of these compounds were -3.2, −3.52, and −3.41 (kcal/mol), respectively (see Fig 8 and supplementary 3).

**Figure 8.**
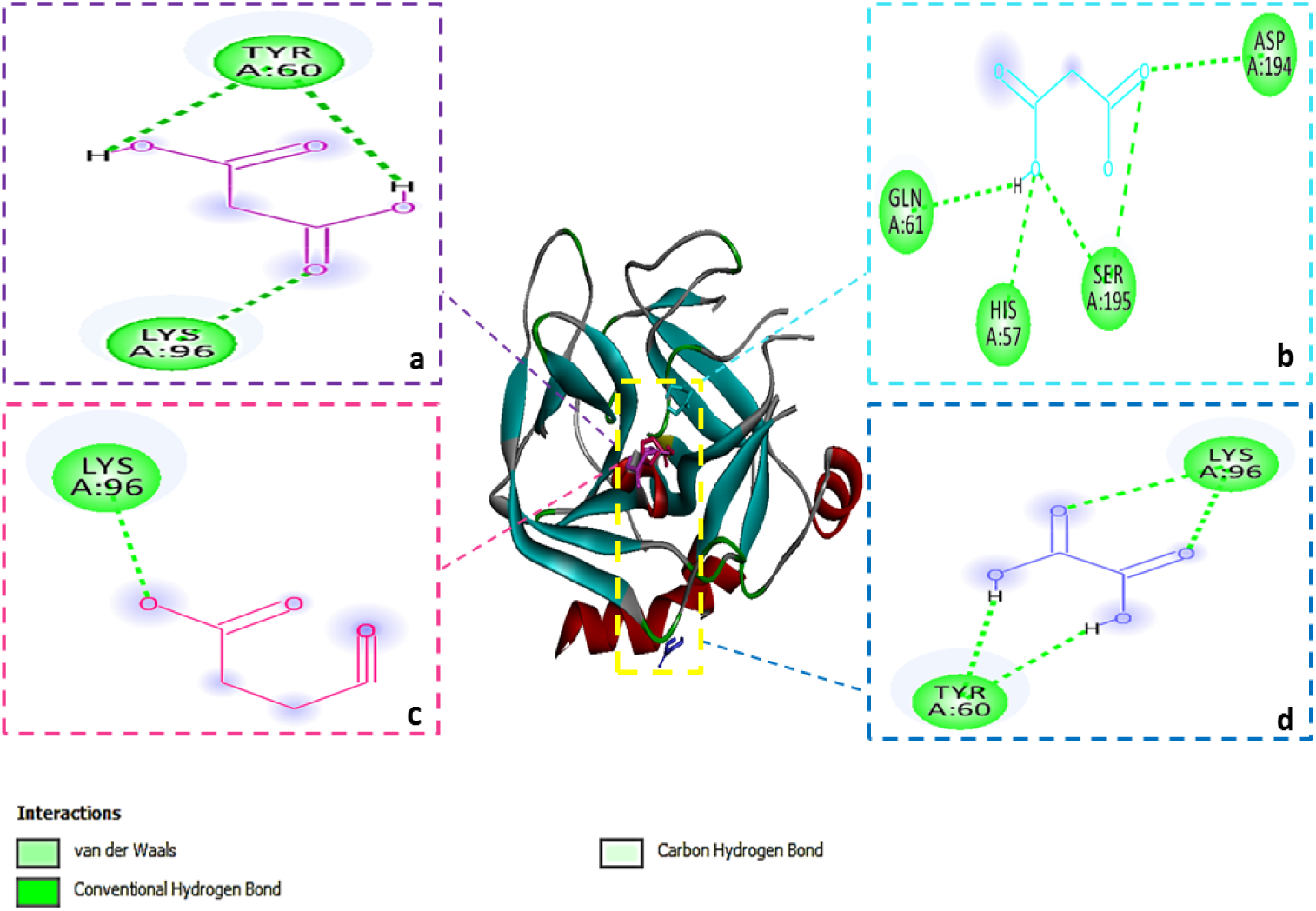
3-D (three dimensional) diagram and 2-D (two dimensional) diagram showing several interactions between malic acid (a), malonic acid (b), succinic acid (c) and oxalic acid (d) and coagulation factor Xa

#### Coagulation FVa

Four investigated ligands interacted with the active site of the protein. The malonic acid had the best docking scores (binding energy -5.01 kcal/mol). It was observed to interact with the receptor through one conventional hydrogen bond with THR-1719 at a distance of 2.19 Å. 3-D and 2-D representations of this compound are shown in Fig 9. The compound malic acid was the second-best ligand (binding energy -4.99 kcal/mol). It was found to interact with the receptor via one conventional hydrogen bond with THR1719 at distances of 2.29 Å (see Fig 9 and supplementary 3).

**Figure 9.**
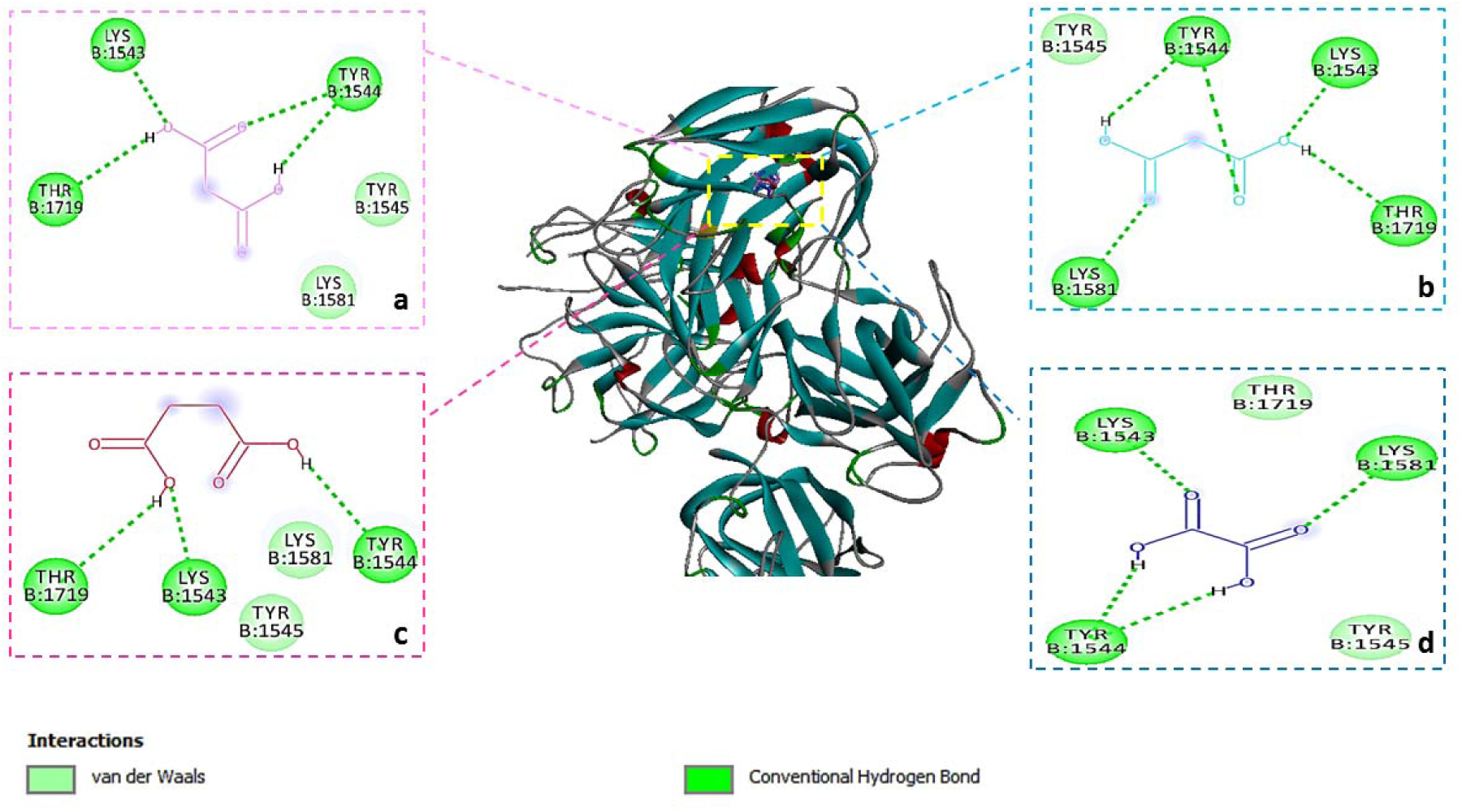
3-D (three dimensional) diagram and 2-D (two dimensional) diagram showing several interactions between malic acid (a), malonic acid (b), succinic acid (c) and oxalic acid (d) and coagulation factor Va

The 3-D and 2-D representations of oxalic acid (binding energy -4.58 kcal/mol) in the active pocket of the coagulation FVa receptor are shown in Fig 9. It is observed to interact with the FVa protein receptor via one van der Waals bond with THR-1719 (see Fig 9 and supplementary 3).

Succinic acid had promising docking scores (-4.51 kcal/mol) (supplementary 3). It is found to interact with the receptor through one van der Waals bond with THR-1719. 3-D and 2-D representations of this compound in complex with the FVa protein receptor are depicted in Fig 9.

#### Coagulation FIIa

During the investigation of the interaction of these compounds with thrombin, all drugs were placed in the active site of this coagulation factor, where the binding energy (-3.19 kcal/mol) and inhibit constant (5.58 millimolar) of malic acid was higher than other compounds (supplementary 3). Malic acid also received the best docking scores (see Fig 10 and supplementary 3). It was observed to interact with the receptor through two conventional hydrogen bonds and a carbon-hydrogen bond with HIS-57 (78) at distances of 1.65Å, and 2.15Å, respectively (Fig 10). Furthermore, this compound interacted with the SER-195 residue in the active site of the protein via a conventional hydrogen bond with a length of 2.66Å. Malonic acid was the second-best ligand (binding energy -3.17 kcal/mol) (supplementary 3). It was found to interact with the receptor via two conventional hydrogen bonds and a carbon-hydrogen bond with HIS-57 at distances of 1.65Å and 2.26Å, respectively (Fig 10). Moreover, this compound formed a conventional hydrogen bond with a length of 2.87Å with SER195 residue. The 3-D and 2-D representations of succinic acid (energy binding -3.07 kcal/mol) in the active pocket of the receptor are shown in Fig 10. It is observed to interact with the active site of protein receptor via two conventional hydrogen bonds with HIS-57 at a distance of 1.71Å and SER-195 at a distance of 2.99Å. Oxalic acid interacted with to the receptor via a conventional hydrogen bond with HIS57 at a distance of 2.64Å and van der Waals link with SER-195 with binding energy of -3 kcal/mol (see Fig10 and supplementary 3). 3-D and 2-D representations of this compound in the active site of the protein receptor are shown in Fig 10. In addition, oxalic acid and succinic acid had van der Waals bonds with SER-195 residue.

**Figure 10.**
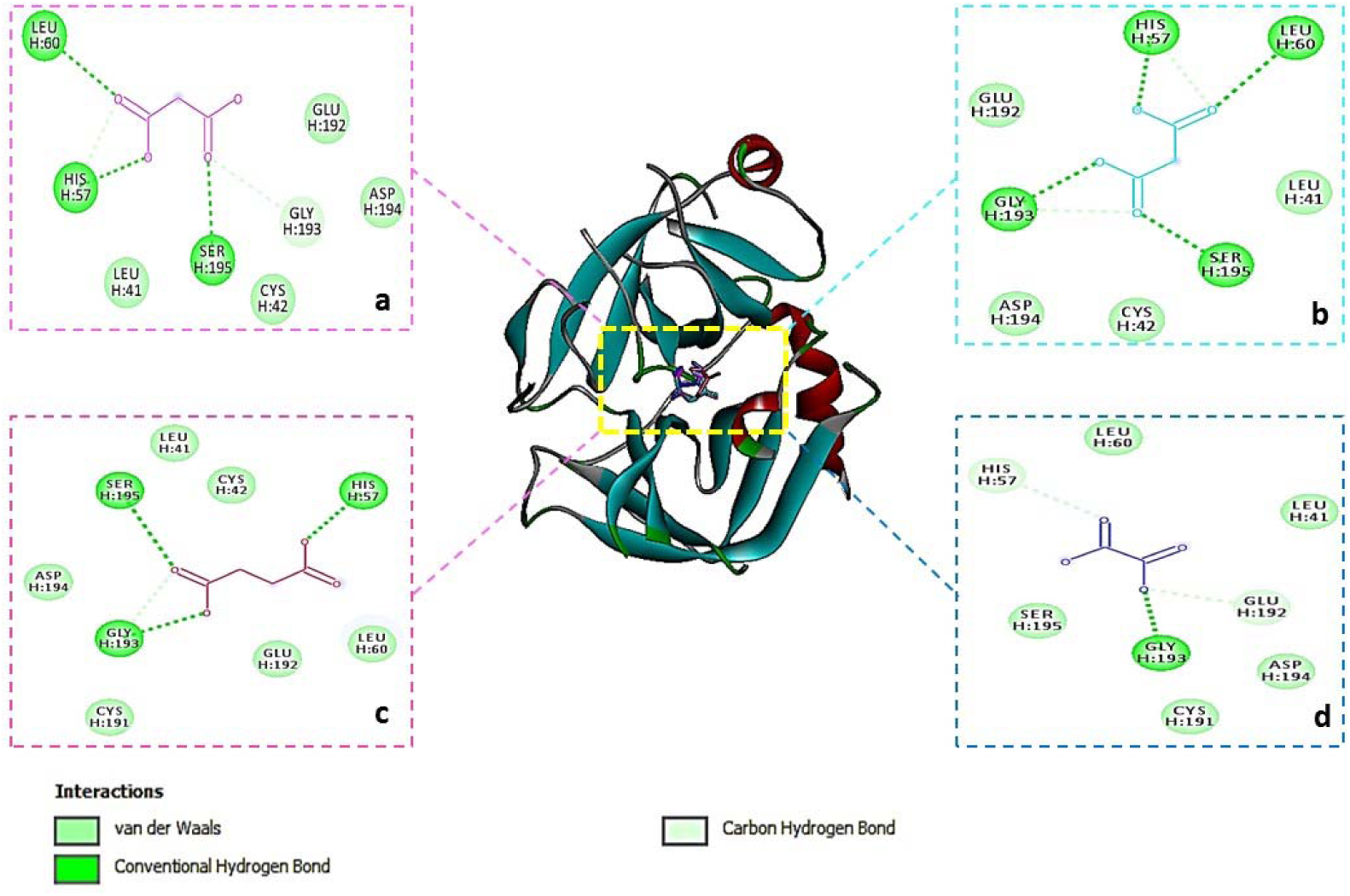
3-D (three dimensional) diagram and 2-D (two dimensional) diagram showing several interactions between malic acid (a), malonic acid (b), succinic acid (c) and oxalic acid (d) and thrombin

#### Coagulation FXIIIa

Malic acid, malonic acid, succinic acid, and oxalic acid interacted with the active site of FXIIIa in the catalytic core with energies of -1.86, -1.69, -1.63, and -1.83 (kcal/mol), respectively. All four ligands interacted with TRP-279 and GLY-277 residues in the active site (79–81) with two van der Waals bonds (see Fig 11 and supplementary 3).

**Figure 11.**
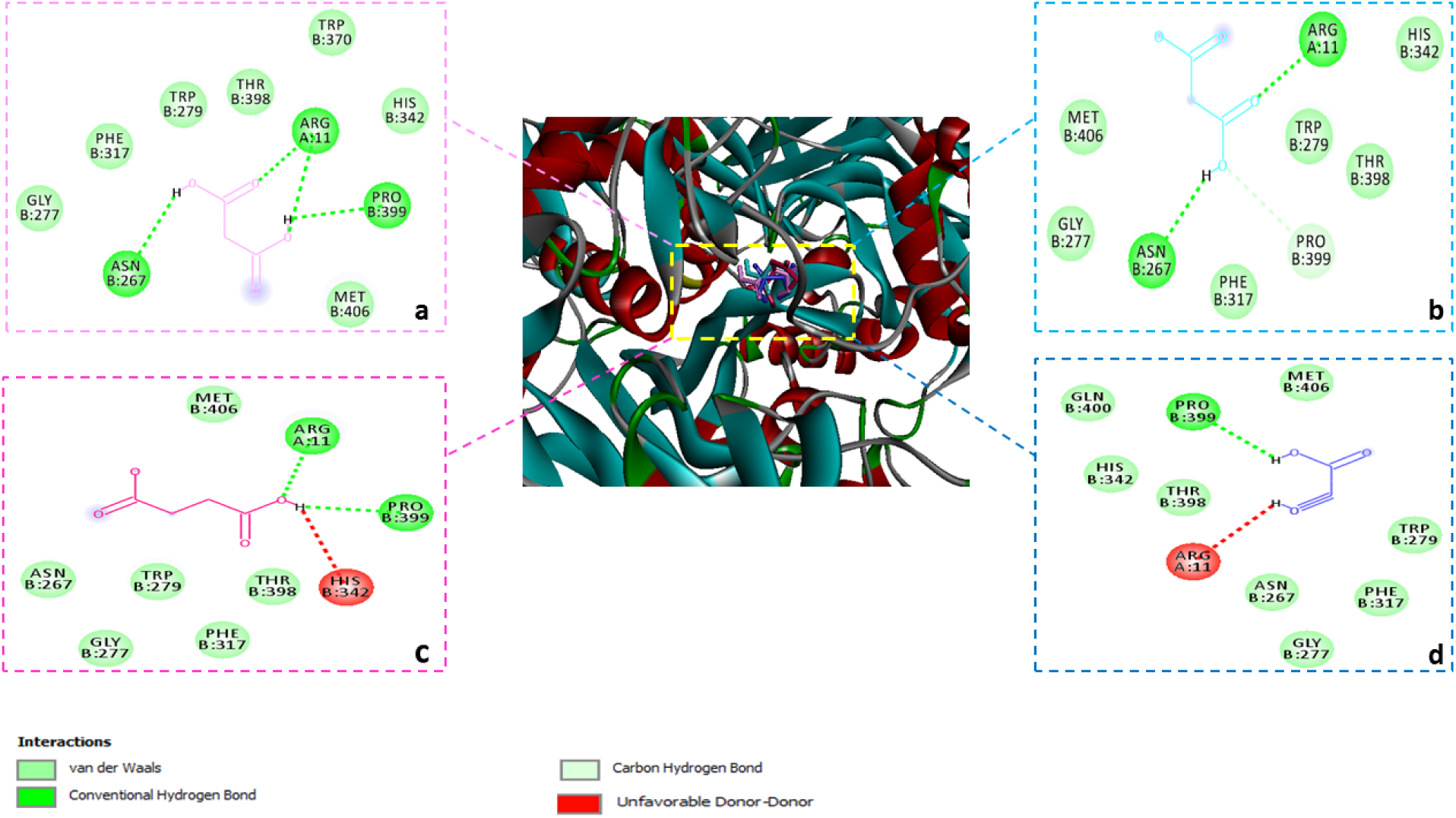
3-D (three dimensional) diagram and 2-D (two dimensional) diagram showing several interactions between malic acid (a), malonic acid (b), succinic acid (c) and oxalic acid (d) and factor XIIIa.

## Discussion

Many therapeutic options have been used for treatment of the thrombosis and hemostasis system disorders, including gene therapy, cell therapy, and replacement therapy, using plasma-derived or recombinant agents. However, these methods may be associated with some disadvantages, such as inhibitor production, short half-life, high drug production cost, cancer risk with gene therapy, and rare transmission of viral infections (14–16, 82–87)

Due to the presence of these side effects, it seems necessary to find more suitable therapeutic options that the body adapts to and stays in the body longer (9, 88–92). The use of effective medicinal plant compounds has played an important role in the treatment of these disorders (36). However, finding herbal compounds to obtain suitable drugs using laboratory methods will be time-consuming and expensive. Therefore, virtual screening, as a computational and validated method, can be used to identify the best and most stable compounds (36). Bioinformatics is an interdisciplinary science that spans drug discovery and uses high-throughput molecular information in comparisons between normal controls and symptom-carriers (93). Molecular docking is an important and appropriate method in the in silico approach that predicts the relationship between a potential ligand and a biological target using a scoring function. Recently, simulation techniques have been used as identification and optimization tools in pharmaceutical programs, fact-based drug target identification, and determining the potential for drug repositioning (94).

Considering that previous studies have shown that 4 compounds present in the C.americanus plant decrease CT, this study aimed to investigate the effect of the identified compounds on the blood coagulation factors. The process of homeostasis prevents bleeding by trapping and keeping blood inside the damaged vessel wall. Hemostasis is a complex process that is divided into primary and secondary hemostasis pathways and depends on the complex interaction of platelets, plasma coagulation cascade, fibrinolytic proteins, blood vessels, and cytokine mediators (95). The internal pathway of blood coagulation includes factors XIIa, XIa, IXa, and VIIIa (96).

To investigate the in silico effect of C.americanus compounds on the intrinsic pathway, all four ligands interacted with the active site of FXIIa, FXIa, and FVIIIa, and the binding energy of malic acid was lower than that of the other compounds. Oxalic acid, in addition to binding to the active site of these factors, interacts with the active site of FIXa. Therefore, it is supposed that these four compounds can alert the function of FXIIa, FXIa, FIXa, and oxalic acid can also affect FVIIIa.

The extrinsic pathway includes tissue factor and factor VIIa (97). FVIIa is a trypsin-like serine protease that forms a complex with an allosteric regulator and tissue factor upon injury and initiates blood clot formation (98). In the prediction of the interaction of C.americanus compounds with FVIIa, all compounds interacted with the active site of this coagulation factor. The binding energy of malic acid was lower than that of the other compounds. Therefore, due to interaction with the active site and lowest binding energy, it binds to FVIIa and activates it.

The common blood coagulation pathway involves the combination of FXa with FV to form the prothrombinase complex, which converts prothrombin to thrombin. Thrombin causes the cleavage of fibrinogen and produces insoluble fibrin, activating FXIII, which covalently connects the fibrin polymers in the platelet plaque. This produces a fibrin network that stabilizes the clot and forms a definitive secondary hemostatic plug (99). In prediction of interaction of C.americanus compounds with the common coagulation cascade pathway, all compounds interacted with the active site of coagulation FVa and FXa, with malonic acid having the lowest energy compared to other compounds in binding to the active site of these two coagulation factors. Furthermore, all the compounds were able to reach the active site of FIIa and FXIIIa, and the binding energy of malic acid was lower than that of other compounds. It seems that these compounds can be effective on most coagulation factors.

It seems that compounds that cause blood coagulation have an aromatic ring, phenolic, ester hydroxyl, carboxylic, and amino group in their structure. Four compounds had two carboxylic and alcoholic groups and appear to interact with coagulation factors with lower energy. In the studies conducted by Muindi et al. to investigate the effects of methanolic extracts of *Croton megalocarpus* and *Lantana camara* plants, the compound quercetin-3-O-B-D-glucoside decreased the activated partial thromboplastin time (APTT) (100, 101). Lin et al. investigated the effects of *Sedum aizoon* compounds on the coagulation system; in this study, Gallic acid (102) caused a reduction in the time of internal blood coagulation in vivo and in vitro, having the same carboxylic acid functional group as our study compounds (102). In the experiment of Yin et al. conducted in 2018 to find blood coagulant compounds in *Malus pumila*, the compound Kaempferol-7-O-β-D-glucopyranoside reduced blood coagulation time of intrinsic pathway and thrombin time (TT). Also, in another study, the composition of vanillic acid in *Sedum aizoon* had a similar effect on the coagulation system. It seems that alcohol and carboxylic acid functional groups in these compounds can cause blood clotting (102, 103). The compounds quercetin-3-O-α-L-rhamnoside, quercetin-3-O-β-D-glucoside, 5-O-coumaroylquinic acid methyl ester, Phloridzin, benzo[3,4]dioxol-10-yl)-7-hydroxypropyl (benzene-2,4-diol), and erythro-2- (4-allyl-2,6-dimethoxyphenoxy)-1-(4-hydroxy-3-rnethoxyphenyl)propan1-ol decreased the clotting time of the extrinsic pathway and TT by having alcohol functional groups (103–105). Also, the alcohol functional group is observed in Pyracanthoside and Kaempferol-3-O-α-L-arabinofuranoside compounds, which reduced blood coagulation time in the external pathway (103).

In the study conducted by Zhang et al. to identify blood coagulant compounds in the Amygdalus persica plant, the compounds 5-O-coumaroylquinic acid, kaempferol-3-O-α-L-rhamnoside, kaempferol-3-O-β-D- galactoside, and D-glucitol were found to decrease TT in vitro. All of these compounds have an alcohol functional group (104). When compared with compounds previously found to be effective in the blood coagulation process, it appears that the functional groups of carboxylic acid and alcohol play an important role in interaction with coagulation factors. In pharmacokinetic assay, all of the compounds exhibited characteristics in agreement with Lipinski’s properties, indicating that they are expected to have no issues with absorption and permeability processes and that they can be considered for further preclinical trials.

Furthermore, all of the compounds had high absorption in MDCK cells, in which, with the exception of oxalic acid, these compounds had the best oral bioavailability and the ability to bind poorly to plasma proteins Our analysis suggests that all compounds correspond to an even distribution between compartments, and all of them have a high fraction unbound in plasma. None of the compounds can cross the BBB penetration, and their transfer is not facilitated by P-gp. None of the compounds inhibit this cytochrome.

None of the compounds inhibit the potassium channels of the heart, cause human hepatotoxicity, liver damage, or oral toxicity in rats. Malic acid and succinic acid are the only compounds that will not cause skin sensitivity (Table 5). In conclusion, carboxylic acid structures and alcoholic groups of the four selected compounds play a role in interacting with coagulation factors. In addition, the compounds’ favorable pharmacokinetic properties make them potential candidates for blood coagulation with fewer side effects in bleeding disorders. Given that previous studies have shown that compounds from this plant reduced CT, and that this study found that these compounds were able to connect with the active site of most coagulation factors with low binding energy, further clinical trials are needed to test their interactions with coagulation factors and accuracy of results. Effective compounds from this plant can be extracted for this purpose.

## Acknowledgments

The authors would like to express their gratitude to Dr. Reza Yakta at the Hong Kong Institute of China for their excellent comments and to Dr. Maryam Sadat Hosseini and Fatemeh Emami for providing a powerful computing system. In addition, the authors would like to thank OpenAI Corporation for providing the free internet platform, GPT 3.5, for editing and improving their manuscript. Finally, the authors also thank the Student Research Committee of Babol University of Medical Sciences for their valuable recommendations.

---

**Supplementary 1.** Interaction of various compounds of C.americanus with intrinsic blood coagulation pathway factors.

| ligand | N subrank | lowet energy(kcal/mol) | Ki (millimolar) | Electrostatic Energy( kcal/mol) | vdW + Hbond + desolv Energy( kcal/mol) |
| --- | --- | --- | --- | --- | --- |
| **Coagulation factor XIIa** |  |  |  |  |  |
| malic acid | 1 | -4.37 | 628.36 | -3.57 | -1.99 |
| malonic acid | 2 | -4.25 | 761.7 | -3.55 | -1.9 |
| succinic acid | 44 | -3.92 | 1.34 | -3.16 | -2.26 |
| oxalic acid | 4 | -3.52 | 2.62 | -3.05 | -1.37 |
| **Coagulation factor XIa** |  |  |  |  |  |
| malic acid | 25 | -3.83 | 1.55 | -2.78 | -2.24 |
| malonic acid | 30 | -3.87 | 1.47 | -2.78 | -2.28 |
| succinic acid | 30 | -3.9 | 1.39 | -3.81 | -1.58 |
| oxalic acid | 48 | -3.74 | 1.81 | -2.48 | -2.16 |
| **Coagulation factor IXa** |  |  |  |  |  |
| malic acid | 24 | -5.57 | 82 | -5.03 | -1.74 |
| malonic acid | 24 | -5.66 | 71.58 | -4.93 | -1.92 |
| succinic acid | 4 | -5.99 | 40.8 | -5.45 | -2.03 |
| Oxalic acid | 4 | -5.15 | 168.46 | -2.98 | -3.06 |
| **Coagulation factor VIIIa** |  |  |  |  |  |
| malic acid | 16 | -2.14 | 26.99 | -2.07 | -1.26 |
| malonic acid | 25 | -2.11 | 28.46 | -1.98 | -1.32 |
| succinic acid | 14 | -1.88 | 42.21 | -2.19 | -1.17 |
| oxalic acid | 12 | -2.52 | 14.2 | -1.87 | -1.54 |

**Supplementary 2.** Interaction of compounds with coagulation factor VIIa.

| ligand | N subrank | lowet energy(kcal/mol) | Ki (millimolar) | Electrostatic Energy( kcal/mol) | vdW + Hbond + desolv Energy( kcal/mol) |
| --- | --- | --- | --- | --- | --- |
| Coagulation factor VIIa |  |  |  |  |  |
| malic acid | 50 | -4.34 | 654.91 | -2.51 | -3.03 |
| malonic acid | 49 | -4.35 | 652.7 | -2.52 | -3.02 |
| succinic acid | 41 | -4.02 | 1.13 | -3.44 | -2.07 |
| Oxalic acid | 50 | -3.55 | 2.49 | -2.53 | -1.91 |

**Supplementary 3.** Interaction of compounds with common blood coagulation pathway.

| ligand | N subrank | lowet energy(kcal/mol) | Ki (millimolar) | Electrostatic Energy( kcal/mol) | vdW + Hbond + desolv Energy( kcal/mol) |
| --- | --- | --- | --- | --- | --- |
| **Coagulation factor Xa** |  |  |  |  |  |
| malic acid | 34 | -3.2 | 4.5 | -2.48 | -1.92 |
| malonic acid | 2 | -3.37 | 3.39 | -2.27 | -2.29 |
| succinic acid | 21 | -3.52 | 2.65 | -3.62 | -1.38 |
| oxalic acid | 48 | -3.41 | 3.18 | -2.44 | -1.86 |
| **Coagulation factor Va** |  |  |  |  |  |
| malic acid | 49 | -4.99 | 220.06 | -3.69 | -2.49 |
| malonic acid | 50 | -5.01 | 213.81 | -3.77 | -2.43 |
| succinic acid | 26 | -4.51 | 497.95 | -3.31 | -2.69 |
| oxalic acid | 50 | -4.58 | 435.77 | -3.06 | -2.42 |
| **thrombin** |  |  |  |  |  |
| malic acid | 41 | -3.19 | 4.58 | -2.33 | -2.05 |
| malonic acid | 46 | -3.17 | 4.71 | -2.32 | -2.04 |
| Succinic acid | 32 | -3.07 | 5.59 | -2.32 | -2.25 |
| Oxalic acid | 50 | -3 | 6.27 | -2.37 | -1.53 |
| **Coagulation factor XIIIa** |  |  |  |  |  |
| malic acid | 32 | -1.86 | 43.1 | -2.28 | -0.77 |
| malonic acid | 23 | -1.69 | 57.29 | -1.79 | -1.09 |
| Succinic acid | 2 | -1.63 | 64.1 | -2.25 | -0.85 |
| Oxalic acid | 50 | -1.83 | 45.82 | -1.68 | -1.04 |

